# Focal adhesion pathway inhibition is the central axis of macrophage phenotypic responses to monoclonal antibody therapy in aggressive lymphoma *via* high-throughput screening and high-content imaging

**DOI:** 10.1101/2025.09.23.677924

**Authors:** Stuart James Blakemore, Peter Zentis, Linda Müller, Reinhild Brinker, Michael Michalik, Christian Jüngst, Anna Beielstein, Jing Zhang, René Neuhaus, Alexandra Florin, Michael Hallek, Jörn Meinel, Reinhard Büttner, Astrid Schauss, Christian Pallasch

## Abstract

High-grade B-cell lymphoma (HGBCL) frequently arises as a refractory or relapsed state of diffuse large B-cell lymphoma (DLBCL) and is associated with poor outcomes due to multi-drug resistance and hallmark oncogenic translocations. To identify novel therapeutic strategies, we developed a dual high-throughput screening (HTS) and high-content imaging (HCI) macrophage–tumour co-culture platform that quantifies antibody-dependent and antibody-independent cellular phagocytosis (ADCP/AICP) across a 1,241-compound library. Using GFP+ HGBCL cells and mCherry+ macrophages, we validated our methodology through time-resolved phenotypic profiling, Euclidean distance-based analysis, and hit compound prioritisation. Pathway interrogation revealed focal adhesion as a central hub of macrophage phenotypic modulation, highlighting focal adhesion kinase (FAK/PTK2) as a candidate therapeutic target. Pharmacological inhibition with PF-562271 enhanced phagocytic activity, altered macrophage morphology, and synergised with anti-CD20 monoclonal antibodies in vitro and ex vivo. In vivo, Rituximab plus PF-562271 significantly reduced lymphoma burden and prolonged survival in xenograft models. Collectively, our work demonstrates that HTS/HCI-driven phenotypic profiling of tumour-associated macrophages can uncover actionable therapeutic combinations and nominates FAK inhibition as a promising strategy to potentiate antibody immunotherapy in HGBCL.

## Introduction

In aggressive B-cell lymphomas, immunochemotherapeutic regimens including the anti-CD20 monoclonal antibody (mAb) Rituximab are curative in ∼60% of Diffuse Large B-Cell Lymphoma (DLBCL) cases^1^, however relapse/refractory cases often acquire multi-drug resistance in the second/third line setting^2^, often gaining *BCL2, MYC*, and/or *BCL6* translocations, defining them as High-Grade B-Cell Lymphoma (HGBCL)^3^. Recently, He *et al*. and Hu *et al*. have offered potential solutions to the identification of novel therapeutic combinations, either *via* high-throughput screening (HTS) luminescence of tumour macrophage co-cultures or high-content imaging (HCI) of M1/M2 polarised macrophage monocultures, respectively^4–6^. Here, we have established a dual HTS/HCI *in vitro* HGBCL-Macrophage co-culture system, leveraging antibody-dependent cellular phagocytosis (ADCP) and antibody-independent cellular phagocytosis (AICP) to characterise tumour associated macrophage (TAM) phenotypic response across a 1,241-compound library, validating the efficacy of this approach using the dual Focal Adhesion Kinase/Protein Tyrosine Kinase 2 (*PTK2/PTK2B*) PF-562271 hit compound *in vivo* and *ex vivo*.

## Results

Whilst in previous work we zoomed in on gene or pathway-specific approaches ^7–9^, our goal here was to develop an ADCP/AICP HTS platform. Using our established human *MYC/BCL2* (hMB) GFP^+^ HGBCL and J774.A1 mCherry^+^ murine macrophage empty vector *in vitro* co-culture system^7–10^, we conducted adherent anti-CD52 mAb Alemtuzumab treated ADCPs *via* suspension cell removal, adherent layer lysis and CD19 MACS separation, followed by flow cytometry (Suppl. Methods). Unexpectedly, we identified significant numbers of GFP^+^/mCherry^+^ singlets in the CD19^-^ fraction under Alemtuzumab treatment (*P*.adj<0.0001, Fig.1A). This suggested that sufficient GFP^+^ signal remained within the macrophages to start developing our screening approach. We tested the stability of this signal, conducting adherent ADCPs either with Alemtuzumab alone or in combination with 10µM of the JAK2 inhibitor Tofacitinib. Not only could we show increased binding of HGBCL cells to macrophages, supporting the therapeutic efficacy of our previous work^7^ (*P*.adj = 0.006, Fig.1B, GFP^+/^ mCherry^-^), but also the stability of the signal (*P*.adj = 0.0992, Fig.1B, GFP^+^/mCherry^+^). Lastly, we optimised our co-culture for 384-well plates and conducted a timeseries experiment for 52-hours (Figure 1C-E). After image segmentation, single cell area, shape, and fluorescence parameters were extracted from mCherry^+^ macrophages using an in-house CellProfiler pipeline, followed by a downstream in-house RStudio analysis script motivated by the work of Young *et al*.^11^ (Figure 1F-H, Suppl. Methods). Briefly, we calculated the main principal factors within the dataset, then conducted the Euclidean distance calculation versus untreated wells specifically for macrophage GFP^+^ as proof of principle. Not only could we observe significant GFP^+^/mCherry^+^ signal within principal factor 2 and 3 of the morphology phenotype radar plots (Fig.1G), but also that GFP^+^ Euclidean distance scores for all 3 mAbs were adequately captured by this analysis, including a significant increase in expression in Rituximab and Rituximab + Tofacitinib versus Tofacitinib monotreatment (*P*.adj<0.1, Fig.1H).

**Figure 1:**
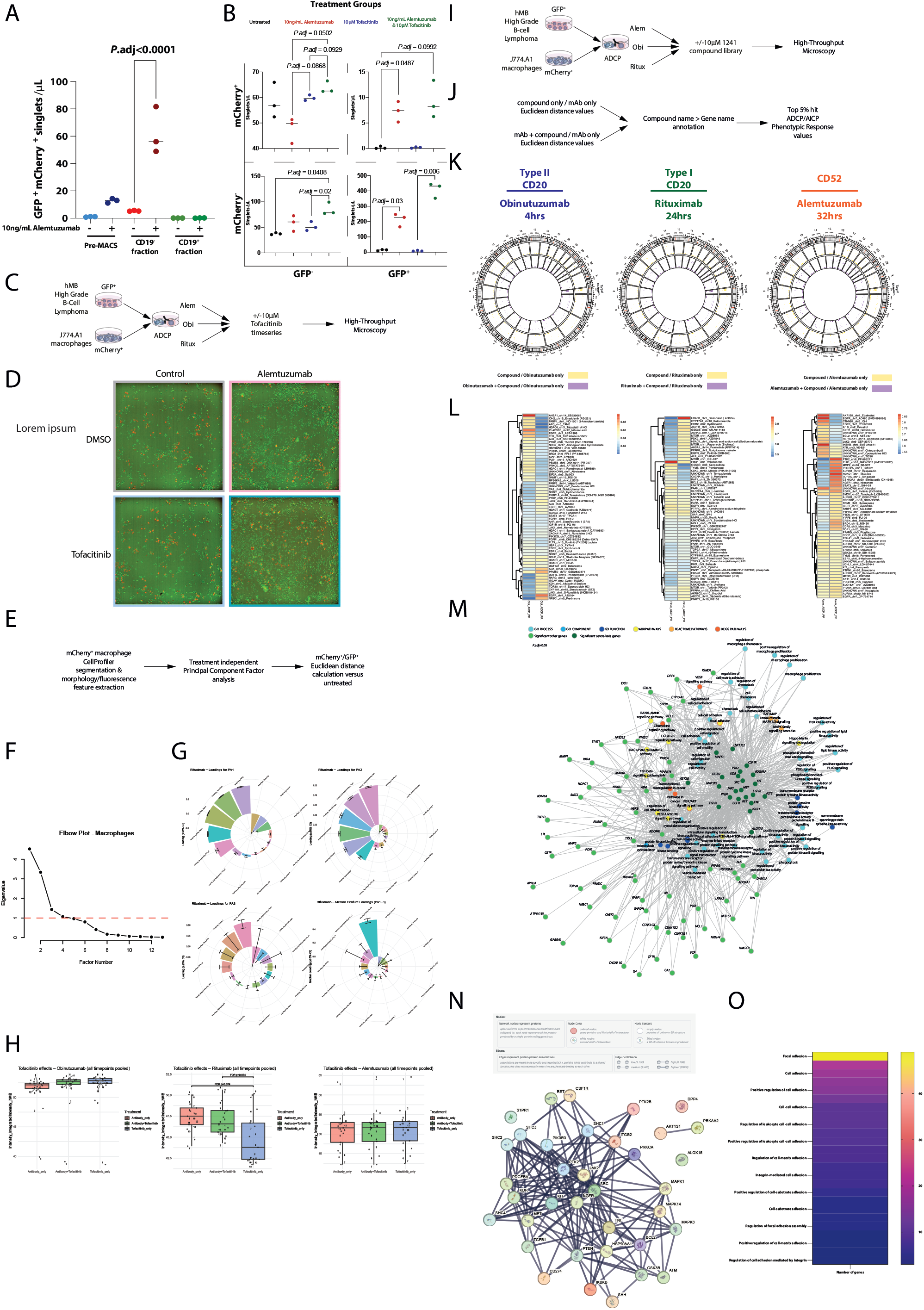
Phenotypic response of tumour associated macrophages unearths focal adhesion as central axis driving HGBCL phagocytosis by high-throughput screening and high-content imaging. **A** Alemtuzumab-mediated upregulation of HGBCL GFP positivity in mCherry^+^ macrophages post CD19+/- MACS separation (3 independent experiments, 3 technical replicates). **B** Adherent Alemtuzumab-mediated upregulation of HGBCL GFP positivity in the context of 10µM Tofacitinib (JAK2) inhibition (3 independent experiments, 3 technical replicates). **C** Schematic overview of the high-content imaging optimisation approach (1 independent experiment per antibody, 96 technical replicates). **D** Representative HGBCL GFP^+^ macrophage mCherry^+^ images under untreated (DMSO), Alemtuzumab, Tofacitinib, or combination treatment. **E** Schematic overview of the analysis pipeline to assess high-content imaging screenability. **F** Elbow plot representing the number of principal factors to include for downstream analysis. **G** Radar plots showing the variability in area shape features within included principal factors. **H** Pooled timeseries bar plots showing Euclidean distance calculation versus untreated cells based on mCherry^+^/GFP^+^ macrophages. **I** Schematic overview of the setup for the high-throughput screening and high-content imaging approach (1 independent experiment per antibody, 3 biological replicates). **J** Schematic overview of the extended analysis pipeline based on median principal factor scores and the inhibitor to gene name annotation approach. **K** CIRCOS plot of the antibody-independent and antibody-dependent cellular phagocytosis phenotypic response values. **L** Heatmaps representing the top 5% of compounds eliciting the highest average AICP/ADCP phenotypic response scores (blue = lowest values, red = highest values). **M** ggnet plot representing the central axis of genes driving associated gene ontology processes (dark green) or peripheral genes (light green). **N** STRING interaction analyses of central axis genes (edge confidence >0.9). **O** Heatmap representing the number of STRING interaction genes associated with adhesion gene ontology pathways (purple = lowest values, yellow = highest values). Significance either represents adjusted *P* values from one-way ANOVA (*P*.adj<0.1), or top 5% of highest phenotypic response values.

We therefore ran 3 HTS/HCI 1,241-compound library screens, selecting each mAb imaging timepoint dependent upon a combination of its known macrophage activation profile as well as our timeseries Tofacitinib Euclidean distances values as a benchmark (Fig.1H&I, ^12,13^). To be able to reliably identify hit compounds, we additionally calculated the normalised Euclidean distance of the phenotypic response variable (Fig.1J). All three timepoints elicited strong upregulation in phenotypic response (Fig.1I, Suppl. Table 1), with 104 genes being included amongst the Top 5% of average ADCP/AICP phenotypic response values (Fig.1.K&L). Pathway analysis revealed a central axis of 22 genetic targets significantly modulating ADCP/AICP phenotypic response, with pathways largely associated with macrophage proliferation, as well as adhesion and motility (Fig.1M), with string interaction analysis unearthing focal adhesion as the hub of this central axis, including the major focal adhesion pathway member Focal Adhesion Kinase (*FAK/PTK2*; Fig.1N&O).

To assess the sturdiness of FAK inhibition on macrophage phenotypic response, we took the FAK inhibitor PF-562271 forward for deep HCI characterisation. Briefly, we conditioned our macrophages either with standard media or HGBCL conditioned media for 72 hours (see Suppl. Methods), following which we seeded our co-culture with a 7-step log2 concentration series (10µM-0.15625µM) alone or in combination with Alemtuzumab, Obinutuzumab, and Rituximab (10ng/mL; Figure 2A). We then imaged the 384-well plate every hour for 48 hours. After identification of principal factors (Figure 2B&C), we observed a persistent increase in phenotypic response across multiple concentrations, both in the AICP and ADCP context of the unconditioned group (Figure 2D; upper left and right). Intriguingly, at first glance, it looks like the effect is greatly diminished in the conditioned group; however, the scale of the polar plots is orders of magnitude larger than in the unconditioned group plots (Figure 2D; lower left and right, Suppl. Fig. 1). To glean an insight into how the macrophage phenotypic response is altered under PF-562271 inhibition, we recalculated the Euclidean distance per biological replicate for each phenotype feature (Suppl. Methods). Crucially, mCherry^+^/GFP^+^ signal with the macrophages was significantly increased under monotreatment in the unconditioned setting (Figure 2E; upper left), whereas mCherry^+^/GFP^+^ signal was significantly improved under conditioned Rituximab PF-562271 inhibitor combination treatment (*P*.adj<0.005, Figure 2E; lower left). Amongst numerous significantly altered phenotypic features, solidity caught our eye since higher solidity scores have been associated with a reduction in protrusion events, an important physiological function of myeloid cells to adhere and migrate around their microenvironment^14^. Without conditioning, the upregulated solidity score showed modest non-significant changes (Figure 2E; upper right), whereas HGBCL conditioning of macrophages conversely drove highly significant upregulation of solidity phenotypic response scores (and therefore reduced protrusion). These results were stable across all antibodies tested (Suppl. Fig. 2-7), and suggest that mAb + PF-562271 combination treatment is forcing the macrophages to be immobile yet highly phagocytically active. To confirm this finding, we reverted to our standard flow cytometry approach, and could show that independent of macrophage conditioning status, significant increases in phagocytosis could be observed in six out of seven concentrations after 16-hours (Unconditioned = *P*.adj<0.05, Conditioned = *P*.adj<0.0001, Figure 2F; upper and lower left), and seven out of seven concentrations after 40-hours (*P*.adj<0.0001, Figure 2F; upper and lower right, Suppl. Fig 8), respectively.

**Figure 2:**
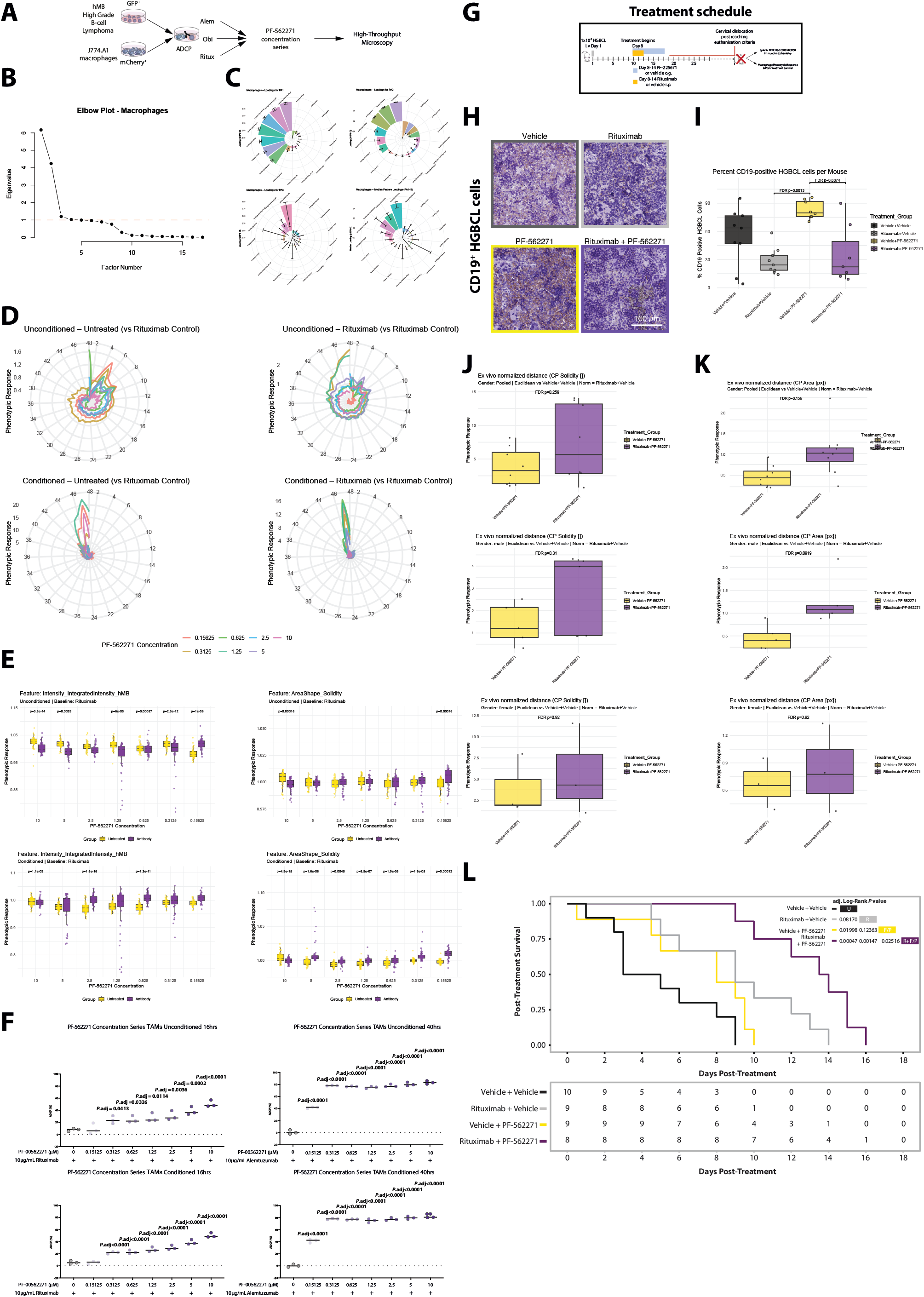
Dual FAK/PYK2 kinase inhibition drives phenotypic response and phagocytosis *in vitro* and *ex vivo*, prolonging post-treatment survival *in vivo*. **A** Schematic overview of the high-content imaging PF-562271 concentration series, timeseries, and prior macrophage conditioning approach (1 independent experiment per antibody, 3 biological replicates). **B** Elbow plot representing the number of principal factors to include for downstream analysis. **C** Radar plots showing the variability in area shape features within included principal factors. **D** Polar plots showing the phenotypic response of each PF-562271 concentration, either ADCP or AICP, with or without conditioning, over time. **E** Pooled timeseries per feature Euclidean distance and phenotypic response calculation of mCherry+/GFP+ (left) or Solidity (right), in the unconditioned (upper) and conditioned (lower) settings (yellow = PF-562271 monotreatment, purple Rituximab + PF-562271 treatment, PF-562271 concentration series [0.15625µM-10µM]). **F** Bar graphs showing the ADCP of Rituximab either alone or in combination with PF-562271 concentrations series for either 16 hours (left) or 40 hours (right), and either with (lower) or without conditioning (upper) (grey = Rituximab monotreatment, shades of purple Ritixumab + PF-562271 concentration series [0.15625µM-10µM]). **G** Schematic overview of the treatment schedule for Rituximab +/- PF-562271 in NSG mice *in vivo*. **H** Representative CD19 DAB expression under untreated (vehicle), Rituximab, PF-562271, or combination treatment. **I** Bar plot representing the percentage of CD19 positive cells per mouse stratified by treatment group (black = vehicle, grey = Rituximab monotherapy, yellow = PF-562271 monotherapy, purple = Rituximab + PF-562271 therapy) (yellow = PF-562271 monotherapy, purple = Rituximab + PF-562271 therapy). **J** Box plots representing phenotypic response scores for Solidity in pooled (upper), male (middle), and female (lower) mice. **K** Box plots representing phenotypic response scores for Area in pooled (upper), male (middle), and female (lower) mice. **L** Kaplan Meier post-treatment survival curve of all mice in the cohort (black = vehicle, grey = Rituximab monotherapy, yellow = PF-562271 monotherapy, purple = Rituximab + PF-562271 therapy). Significance either represents adjusted *P* values from one-way ANOVA (*P*.adj<0.1), or adjusted Log-Rank *P* values (*P*.*adj*<0.1).

Finally, to validate the HCI methodology and assess the pre-clinical efficacy of combined Rituximab + PF-562271 inhibition *in vivo*, we injected HGBCL cells *i*.*v*. into NSG mice, allowed them to engraft for 8-days, following which we initiated Rituximab therapy once daily on day 8, 9, and 10 i.*p*., as well as PF-562271 inhibitor treatment twice daily from day 8-14 *o*.*g*. (Figure 2G; Suppl. Methods). We then monitored the mice until they reached euthanasia criteria, upon which we harvested the spleens for fixation and preparation for immunohistochemistry of human CD19 and murine CD68 antibody staining (Suppl. Methods). Representative CD19 staining of the spleens of these mice stratified by treatment showed significant lymphoma infiltration under vehicle and PF-562271 monotreatment (Figure 2H; upper and lower left), whilst minimal CD19 staining could be observed in Rituximab and Rituximab + PF-562271 combination treatment (Figure 2H; upper and lower right). After cell segmentation of all *ex vivo* splenic samples, calculation of the proportion of CD19 positive cells stratified by treatment group confirmed our visual observations, showing significant reductions in tumour cells under both Rituximab and Rituximab + PF-562271 treatment (*P*.adj<0.01, Figure 2I). Expanding our analysis to the CD68+ macrophages, although an expanded number of principal factors could be defined (Suppl. Fig 9), only sex-stratified analyses restricted to male mice revealed phenotype features that were significant after false discovery rate correction. Regardless, not only could we see non-significant increases in solidity in Rituximab + PF-562271 treated mice in both pooled and stratified by gender (Figure 2J) but also significant increase in Area (*P*.adj = 0.09, Figure 2K), Perimeter, and Major Axis Length (*P*.adj = 0.0275 & 0.0596, Suppl. Fig 10-12) in macrophages of male mice, all of which have been previously reported to correlate with focal adhesion leading or rear edge dynamics^14^. Crucially, we report that Rituximab + PF-562271 not only leads to significantly prolonged post-treatment survival versus all other treatment arms (Log-Rank *P*.adj<0.05, Figure 2L), but that the effect upon survival occurs in a gender independent manner (*P*.adj<0.1, Suppl. Fig 13).

## Discussion/Conclusion

Using our HTS/HCI approach, we show the efficacy of focusing on macrophage phenotypic response to identify novel therapeutic combinations *in vitro*, which lead to improved survival *in vivo*. Many of the inhibitors in the central axis are either clinically licensed or currently undergoing clinical trials, including for FAK inhibitors across various cancers ^15^, supporting the validity of our methodology. PF-562271 itself has currently only been tested in one phase 1 clinical trial in a diverse selection of advanced solid tumours including prostate cancer, but its study was spurious in its design, attempting to associate its toxicity profile with dietary restrictions, and was only tested as a monotherapy^16,17^. Pharmacological inhibition of members of the focal adhesion or its associated pathways is often implicated in pre-clinical studies, frequently without explicit implication of its adhesion-related impact on the TME. Although outside the scope of the current work, we foresee a strategy where this approach could be instigated in the *ex vivo* context of HGBCL patient biopsies to provide novel off-label therapeutic options for these patients with dismal prognoses. Regardless, leveraging HTS/HCI to assess macrophage phenotypic response is another tool in the arsenal to tackle HGBCL.

## Acknowledgements

This study was supported by: the Deutsche Forschungsgemeinschaft (German Research Foundation) grant 455784452 as part of CRC1530 (with project funding for M.H. (B01, Z01), R.B. (Z02), C.P. (B02), the José Carreras Leukämie Stiftung (José Carreras Leukaemia Foundation) grant DJCLS 07R_2021 and DJCLS 07R/2025, the Köln Fortune Programm (Cologne Young Scientists Program) grant 454/2020 and 199/2022 S.J.B, the Exzellenz initiieren Stiftung Kölner Krebsforschung (Initiating excellence Cologne Cancer Research Foundation), S.J.B. All animal experiments were performed according to ethics obtained from Landesamt für Verbraucherschutz und Ernährung (LAVE; State Office for Consumer Protection and Food, Düsseldorf, Germany) nr. 81-02.04.2021.A242.

## Author Contributions

S.J.B, and C.P designed the study, performed experiments, analysed data, and wrote the manuscript. P.Z analysed data. L.M, R.B^1^, M.M, A.B, J.Z, R.N, A.F performed experiments. C.J, M.H, J.M, R.B^2^, and A.S provided resources. All authors read, revised and approved the manuscript.

## References

1. Coiffier, B. et al. Long-term outcome of patients in the LNH-98.5 trial, the first randomized study comparing rituximab-CHOP to standard CHOP chemotherapy in DLBCL patients: a study by the Groupe d’Etudes des Lymphomes de l’Adulte. Blood 116, 2040–2045 (2010).

2. Neelapu, S. S. et al. Five-year follow-up of ZUMA-1 supports the curative potential of axicabtagene ciloleucel in refractory large B-cell lymphoma. Blood 141, 2307–2315 (2023).

3. Alaggio, R. et al. The 5th edition of the World Health Organization Classification of Haematolymphoid Tumours: Lymphoid Neoplasms. Leukemia vol. 36 1720–1748 Preprint at 10.1038/s41375-022-01620-2 (2022).

4. Cao, X. et al. Targeting macrophages for enhancing CD47 blockade–elicited lymphoma clearance and overcoming tumor-induced immunosuppression. Blood 139, 3290–3302 (2022).

5. He, Z., Hu, Z., Wang, L., Xiao, Y. & Cao, X. A protocol for high-throughput screening for small chemicals promoting macrophage-mediated tumor cell phagocytosis in mice. STAR Protoc 6, (2025).

6. Hu, G. et al. High-throughput phenotypic screen and transcriptional analysis identify new compounds and targets for macrophage reprogramming. Nat Commun 12, (2021).

7. Barbarino, V. et al. Macrophage-mediated antibody dependent effector function in aggressive B-cell lymphoma treatment is enhanced by ibrutinib via inhibition of JAK2. Cancers (Basel) 12, 1–25 (2020).

8. Beielstein, A. C. et al. Macrophages are activated toward phagocytic lymphoma cell clearance by pentose phosphate pathway inhibition. Cell Rep Med 5, (2024).

9. Izquierdo, E. et al. Extracellular vesicles and PD-L1 suppress macrophages, inducing therapy resistance in TP53-deficient B-cell malignancies. Blood 139, 3617–3629 (2022).

10. Pallasch, C. P. et al. Sensitizing protective tumor microenvironments to antibody-mediated therapy. Cell 156, 590–602 (2014).

11. Young, D. W. et al. Integrating high-content screening and ligand-target prediction to identify mechanism of action. Nat Chem Biol 4, 59–68 (2008).

12. Stanglmaier, M., Reis, S. & Hallek, M. Rituximab and alemtuzumab induce a nonclassic, caspase-independent apoptotic pathway in B-lymphoid cell lines and in chronic lymphocytic leukemia cells. Ann Hematol 83, 634–645 (2004).

13. Herter, S. et al. Preclinical activity of the type II CD20 antibody GA101 (obinutuzumab) compared with rituximab and ofatumumab in vitro and in xenograft models. Mol Cancer Ther 12, 2031–2042 (2013).

14. Holz, D. & Vavylonis, D. Building a dendritic actin filament network branch by branch: models of filament orientation pattern and force generation in lamellipodia. Biophysical Reviews vol. 10 1577–1585 Preprint at 10.1007/s12551-018-0475-7 (2018).

15. Yang, M., Xiang, H. & Luo, G. Targeting focal adhesion kinase (FAK) for cancer therapy: FAK inhibitors, FAK-based dual-target inhibitors and PROTAC degraders. Biochemical Pharmacology vol. 224 Preprint at 10.1016/j.bcp.2024.116246 (2024).

16. Roberts, W. G. et al. Antitumor activity and pharmacology of a selective focal adhesion kinase inhibitor, PF-562,271. Cancer Res 68, 1935–1944 (2008).

17. Infante, J. R. et al. Safety, pharmacokinetic, and pharmacodynamic phase I dose-escalation trial of PF-00562271, an inhibitor of focal adhesion kinase, in advanced solid tumors. Journal of Clinical Oncology 30, 1527–1533 (2012).

