## Supplemental Methods for "Focal adhesion pathway inhibition is the central axis of macrophage phenotypic responses to monoclonal antibody therapy in aggressive lymphoma *via* high-throughput screening and high-content imaging"

### Cell lines

The murine macrophage cell line J774A.1 and the human HEK 293T were cultured in DMEM, supplemented with 10% FBS and 1% P/S. The human- MYC/BCL2 (hMB) HGBCL cell line (strain 102) was generated by Leskov et al[1], and were cultured in 50% IMDM and 50% RPMI, supplemented with 10% FBS, 1% P/S, and 0.1% 2-Mercaptoethanol.

### Transduction

An MLS-mCherry empty vector was used to generate mCherry$^+$ J774A.1 macrophages by retroviral transduction. Here, HEK 293T derived amphotrophic phoenix cells were transfected by calcium phosphate method with 25$\mu$M chloroquine of an MLS-mCherry empty vector. After 48 h of incubation the virus containing medium was harvested, filtered through a 0.45 $\mu$m syringe filter and integrated by spin infection (400 g 32°C 45 min) on the J774A.1 macrophages. Then, macrophages were incubated with polybrene for 48 h at 37°C before antibiotic selection undertaken with 3 $\mu$g/ml puromycin (Invivo Gen, San Diego, CA, USA) for at least 1 week prior to FACS sorting.

### Antibody-Dependent and -Independent Cellular Phagocytosis flow cytometry assay (ADCP/AICP)

ADCP were performed as described previously [2–4]. Briefly, 1x10$^5$/mL mCherry-positive J77A4.1 macrophages were co-cultured after attachment with HGBCL cells (1.5x10$^6$ cells/mL) in combination or not with the corresponding monoclonal antibody. The anti-CD52 antibody Alemtuzumab (Genzyme, Cambridge, MA, USA), the anti-CD20 Type I antibody Rituximab (Genentech, Inc, South San Francisco, CA and IDEC Pharmaceutical Corporation, San Diego, CA, USA), and the anti-CD20 Type II antibody Obinutuzumab (GlychArt Biotechnology, Zürich, Switzerland) were used at 10µg/ml. Each condition was performed with five replicates in 96-well plates. Remaining GFP+ target cells (hMB HGBCL cells) were analysed by flow cytometry. The percentage of ADCP was calculated as follows: 100 - (100 x (cells/$\mu$L treated / cells/$\mu$L untreated)).

### Adherent ADCP/AICP flow cytometry assay

Using the same plating strategy as above, macrophages and HGBCL cells were seeded in 6-well plates for 16 hours, either with PBS, 10$\mu$g/mL Alemtuzumab or 10$\mu$M Tofacitinib (Pfizer, New York City, USA). After reaching the experimental endpoint, the suspension fraction was removed, and the adherent fraction was washed three times with 3mL MACS buffer, including swirling of the plate and multiple resuspending of the buffer from the edge of the wells. Following which, 1.5mL TryPLE was added to the wells, the plate was briefly returned to the incubator, checking once per minute and with swirling of the plate to confirm effective dissociation from the plate. After this, the reaction was stopped with 1.5mL MACS buffer, transferred to FACS tubes and centrifuged for 5 minutes at 1200rpm. Cells we then either measured directly on an MACSQuant VYB flow cytometer (Miltenyi Biotec), or prior to which underwent human CD19$^+$ MACS bead positive selection. mCherry/GFP negativity/positivity was calculated after gating compensation for the Y2 mCherry/Texas Red and B1 FITC/GFP channels, with downstream analysis assessing significant differences in cells/$\mu$L between treatment groups.

### Generation of conditioned media

1.5x10$^6$/mL HGBCL cells were incubated without treatment for 48-hours. Afterwards, cells were washed and the generated supernatant was collected and centrifuged for 5 min at 350g, filtered through a 0.2$\mu$m sieve and stored at -80°C for flow cytometry and imaging experiments.

**High-Throughput Screening and High-Content Imaging seeding assay**

For the HTS/HCI ADCP/AICP library screening, mCherry+ J774A.1 macrophages were dispensed on 384-well plates in a concentration of 2.5 x $10^3$ cells per well using the MultiFlo FX multi-mode dispenser (BioTek). At least 4 hours later, the GFP+ hMB HGBCL cells were added to the plates in a concentration of 3.75 x $10^4$ cells per well. In total 24 384-well plates were used: 1241 compounds on four 384-well plates (including control wells only containing DMSO), three replicates per compound plate, either alone or in combination with Alemtuzumab, Obinutuzumab, or Rituximab at 10ng/mL. After the hMB cell were dispensed on the plates, 10 $\mu$M compounds were added, using the CyBio FeliX automated pipetting robot (Analytik Jena, Germany). Subsequently, 10 ng/ml alemtuzumab or BCM medium was added to the plates. The plates were measured using the ImageXpress Micro4 (Molecular Devices, USA) high-throughput microscope after optimised timepoints, respectively, capturing images in the brightfield, FITC, and TRITC channels at 10X magnification.

**High-Throughput Screening and High-Content Imaging pre-processing**

Images were pre-processed using an optimised in house CellProfiler pipeline. After loading the appropriate image files, metadata is extracted based upon the folder structure and folder nomenclature. Following which, the NamesAndTypes module was used to name of the FITC and TRITC channells, assigning them to hMB HGBCL and J774A.1, respectively. Next, groups based upon the associated well identifier and group number based on fluorescence channel are optimised, including their location path. After which, speckles were enhanced specifically in J774A.1 cells with a feature size of 35, based upon knowledge of general size of the cells in the cell line. Next, we assign the typical minimum and maximum diameter of the objects (15, 50), as well as a number of other parameters to control for false-positive segmentation of HGBCL cells (such as lower and upper bounds of threshold, smoothing scale, and correction factor). After which, measure object intensity and size and shape parameters are selected for J774A.1 cells. Finally, export information was filled out so that after pre-processing downstream analysis could be conducted.

**High-Throughput Screening and High-Content Imaging analysis**

The pre-processed CellProfiler Macrophage and Image .csv files were firstly used to annotate treatment specific metadata to create Identifier .csv files using Microsoft Excel. Then, all files were loaded into RStudio ready for downstream analysis (experiment specific scripts can be found on GitHub ()). Briefly, data is firstly further pre-processed so that Identifier metadata is correctly annotated onto the Image and Macrophage data frames. Following which, principal factor analysis is conducted independent of treatment groups, excluding phenotypic features which have a correlation R squared value of >0.95. Elbow plots were employed to assess the number of principal factors to include in downstream analysis. Radar plots were generated to define which features group within which factors of the data. The median of included principal components was taken forward to assign treatment specific changes in macrophage morphology, calculating the Euclidean distance between downsampled treated and untreated cells was conducted across all wells belonging to the same treatment group, since the DMSO treatment wells was X in comparison to n = 3 treatment wells per drug/antibody combination. To resolve ADCP and AICP specific modulations in morphology in relation to antibody, normalised Euclidean distance values were conducted by dividing compound monotreatment (AICP) or combination treatment (ADCP) Euclidean distance values by antibody monotreatment, creating Phenotypic Response values. Compounds were considered significant if they were reported in the top 5% of either the AICP or ADCP setting. *In vitro* PF-562271 dual concentration series and timeseries datasets with Alemtuzumab, Obinutuzumab, as well as Rituximab, and the *ex vivo* Rituximab +/- PF-225672 dataset were analysed differently to the main screen. After standard principal factor, Euclidean distance, and

Phenotypic Response calculation, the same analytical framework was repeated for *in vitro* datasets but on a per well within each treatment group basis, so that all AreaShape features and principal factors could be tested for significance by one-way anova. For the *ex vivo* dataset, human CD19+ and murine CD68 expression cutoffs were set independent of sample, treatment group, or gender. Due to the lack of technical replication, only principal factor analysis was conducted on the cohort level, with Euclidean distance and Phenotypic Response being calculated for all available features and principal factors on pooled, gender stratified, and treatment group basis.

**The hMB humanized High-Grade B-Cell Lymphoma model**

Human hematopoietic stem cells were isolated from healthy donor cord blood using $CD133^{+}$ selection and expanded *in vitro*. After infection with the lentiviral CD19-promotor/Eμ-enhancer GFP-MYC-BCL2-construct, cells were injected into sublethally irradiated NOD-scid Il2rg−/− (NSG) mice and monitored for disease onset by counting the number of GFP+ cells in the peripheral blood. Secondary NSG recipient mice were injected with $1x10^{6}$ cells derived from spleens of primary leukemic mice. The hMB model accurately recapitulates the histopathological and clinical aspects of steroid-, chemotherapy- and rituximab-resistant human "High-Grade B-Cell" lymphomas that involve the MYC and BCL2 loci. Here, 8-14 weeks old male NOD.Cg-PrkdcscidIl2rgtm1Wjl/SzJ (NSG, Jackson Laboratory, USA) immunodeficient mice were injected *i.v.* with $1x10^{6}$ hMB HGBCL cells diluted in 100 *μ*l PBS. Unless otherwise stated, 8 days after injection mice were treated *i.p.* on three consecutive days with Rituximab (day 8: 2 mg/kg, day 9 and day 10: 7 mg/kg). Dual FAK/PYK2 inhibitor PF-562271 (Verastem Inc., Massachusetts, United States) was dosed at 33 mg/kg in 100 *μ*l 5% Gelucire 44/14, with a total of 14 administrations *via* oral gavage; 7 consecutive days starting on day 8; twice daily due to the short half-life of 6 hours. PBS *i.p.* and 5% Gelucire *o.g.* were the vehilches for Rituximab and PF-562271, respectively. Disease progression was monitored by weekly blood sampling and daily scoring of the mice. NSG immune incompetent mouse models have been reported to show defective macrophage function. However, in our context of using humanized HGBCL cell lines *in vivo* as well as human therapeutic mAb treatment we have observed functional phagocytic capacity both with chemotherapeutics[5] as well as under numerous small molecule inhibitors[2–4]. Therefore, we believe that this model represents a novel strategy to assess phagocytic capacity using frontline novel targeted therapeutics.

**Ex vivo spleen harvesting, fixation, and imaging**

After preparation of survival cohort of hMB HGBCL transfected NSG mice after treatment with Rituximab and/or PF-562271, a 2 3 2mm piece of spleen was fixated in formaldehyde. Histological sections were produced and immunohistochemical staining of CD19 and CD68 was performed. Prepared splenic sections were imaged on a NanoZoomer S360 (Hamamatsu Photonics, Hamamatsu, Japan) under 400x magnification (0.2305 *μ*m/px). Following which, images were segmented and pre-processed in QuPath, adapting AreaShape parameters and definitions from CellProfiler.

**Statistics**

Data were analysed using GraphPad Prism 8.0 Prism software (San Diego, CA, USA), and R. Results are expressed as median ± SD and medians were compared by Student's t-test, Mann-Whitney U-test, and Wilcoxon tests as appropriate. Statistical comparison between groups was performed using one-way ANOVA multiple comparison test in the ADCP assays. Kaplan Meier survival analysis was performed using pairwise Log-rank (Mantel-Cox) test. Differences were considered statistically significant at p-values less than 0.05 (*, $p \leq 0.05$; **, $p \leq 0.01$; ***, $p \leq 0.001$; ****, $p \leq 0.001$).

1. Leskov, I. *et al.* Rapid generation of human B-cell lymphomas via combined expression of Myc and Bcl2 and their use as a preclinical model for biological therapies. *Oncogene* 32, 1066–1072 (2013).
2. Barbarino, V. *et al.* Macrophage-mediated antibody dependent effector function in aggressive B-cell lymphoma treatment is enhanced by ibrutinib via inhibition of JAK2. *Cancers (Basel)* 12, 1–25 (2020).
3. Beielstein, A. C. *et al.* Macrophages are activated toward phagocytic lymphoma cell clearance by pentose phosphate pathway inhibition. *Cell Rep Med* 5, (2024).
4. Izquierdo, E. *et al.* Extracellular vesicles and PD-L1 suppress macrophages, inducing therapy resistance in TP53-deficient B-cell malignancies. *Blood* 139, 3617–3629 (2022).
5. Pallasch, C. P. *et al.* Sensitizing protective tumor microenvironments to antibody-mediated therapy. *Cell* 156, 590–602 (2014).
