## Supplemental Figures for "Focal adhesion pathway inhibition is the central axis of macrophage phenotypic responses to monoclonal antibody therapy in aggressive lymphoma *via* high-throughput screening and high-content imaging"

**Supplemental Figure 1: Alemtuzumab & Obinutuzumab timeseries analyses of PF-562271 concentration series in the context of macrophage conditioning**

**A** Polar plots showing the phenotypic response of each PF-562271 concentration, either ADCP or AICP, with or without conditioning, over time. **B** Pooled timeseries per feature Euclidean distance and phenotypic response calculation of mCherry+/GFP+ (left) or Solidity (right), in the unconditioned (upper) and conditioned (lower) settings (yellow = PF-562271 monotreatment, purple Alemtuzumab + PF-562271 treatment, PF-562271 concentration series [0.15625$\mu$M-10$\mu$M]). **C** Polar plots showing the phenotypic response of each PF-562271 concentration, either ADCP or AICP, with or without conditioning, over time. **D** Pooled timeseries per feature Euclidean distance and phenotypic response calculation of mCherry+/GFP+ (left) or Solidity (right), in the unconditioned (upper) and conditioned (lower) settings (yellow = PF-562271 monotreatment, purple Obinutuzumab + PF-562271 treatment, PF-562271 concentration series [0.15625$\mu$M-10$\mu$M]).

**A**

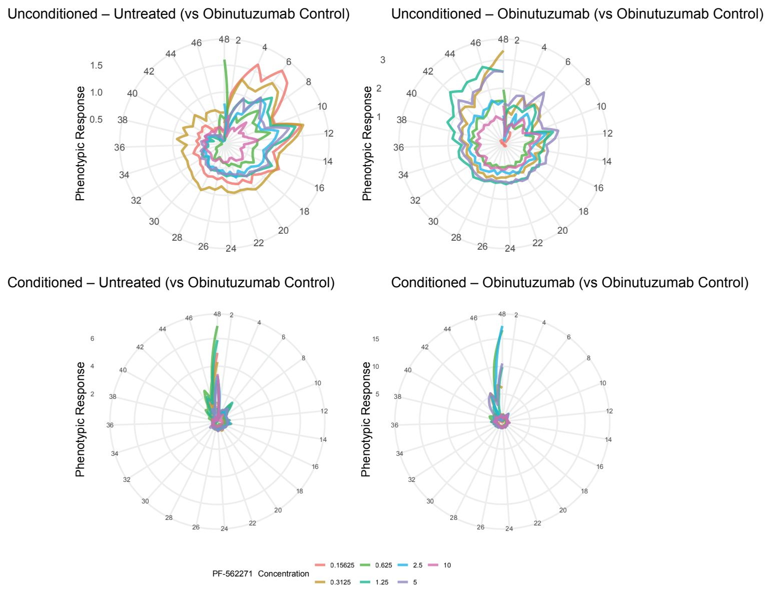


**B**

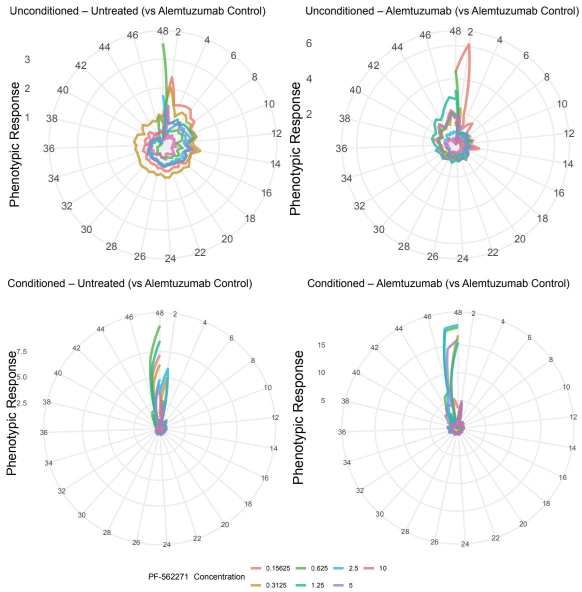

**Supplemental Figure 2: Rituximab timeseries of PF-562271 with per feature phenotypic response calculations without conditioning**

Pooled timeseries per feature Euclidean distance and phenotypic response calculation of all features included in the analysis (n = 24), in the unconditioned setting (yellow = PF-562271 monotreatment, purple Rituximab + PF-562271 treatment, PF-562271 concentration series [0.15625$\mu$M-10$\mu$M]).

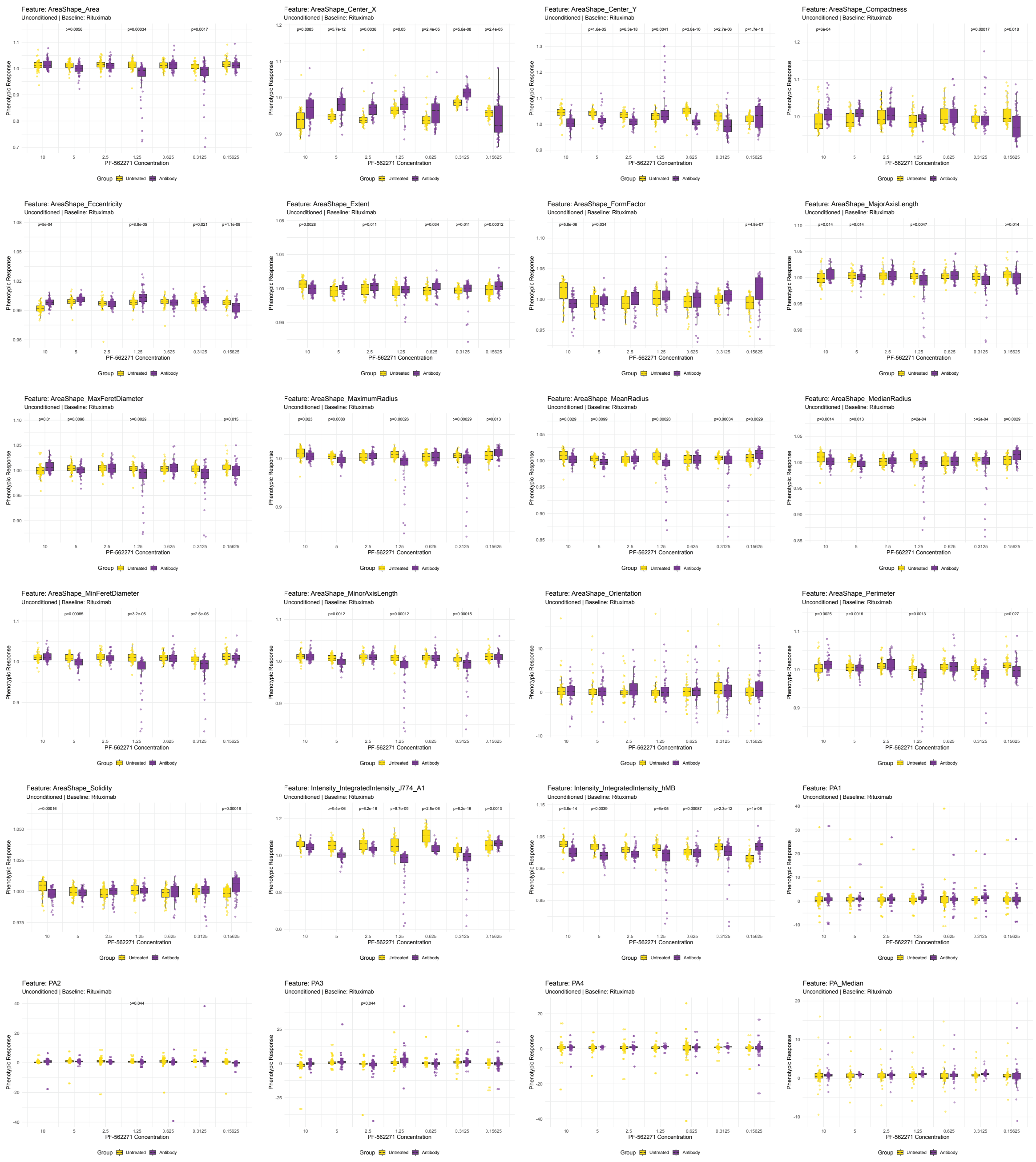

### Supplemental Figure 3: Rituximab timeseries of PF-562271 with per feature phenotypic response calculations with conditioning

Pooled timeseries per feature Euclidean distance and phenotypic response calculation of all features included in the analysis (n = 24), in the conditioned setting (yellow = PF-562271 monotreatment, purple Rituximab + PF-562271 treatment, PF-562271 concentration series [0.15625$\mu$M-10$\mu$M]).

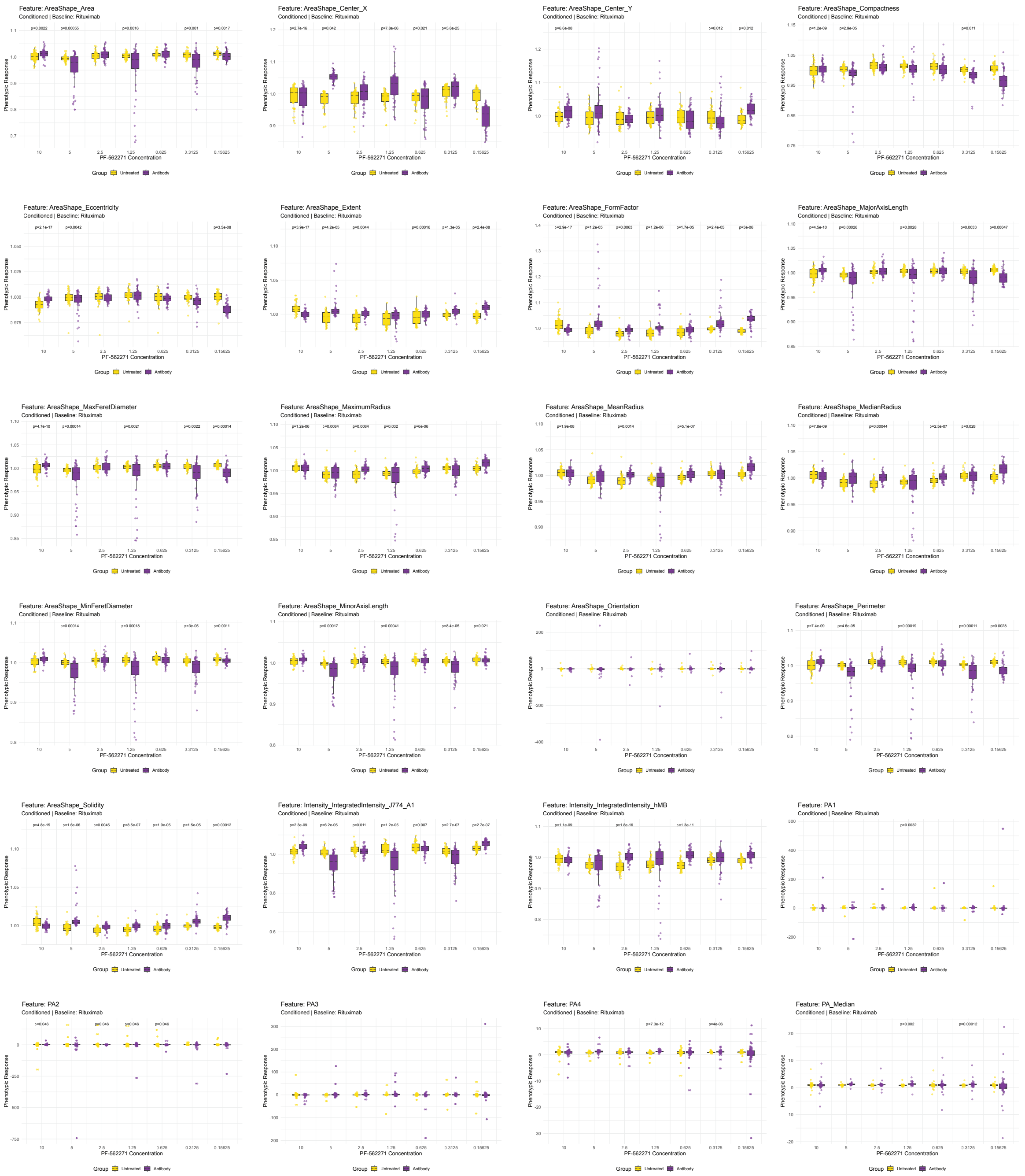

**Supplemental Figure 4: Obinutuzumab timeseries of PF-562271 with per feature phenotypic response calculations without conditioning**

Pooled timeseries per feature Euclidean distance and phenotypic response calculation of all features included in the analysis (n = 24), in the unconditioned setting (yellow = PF-562271 monotreatment, purple Obinutuzumab + PF-562271 treatment, PF-562271 concentration series [0.15625$\mu$M-10$\mu$M]).

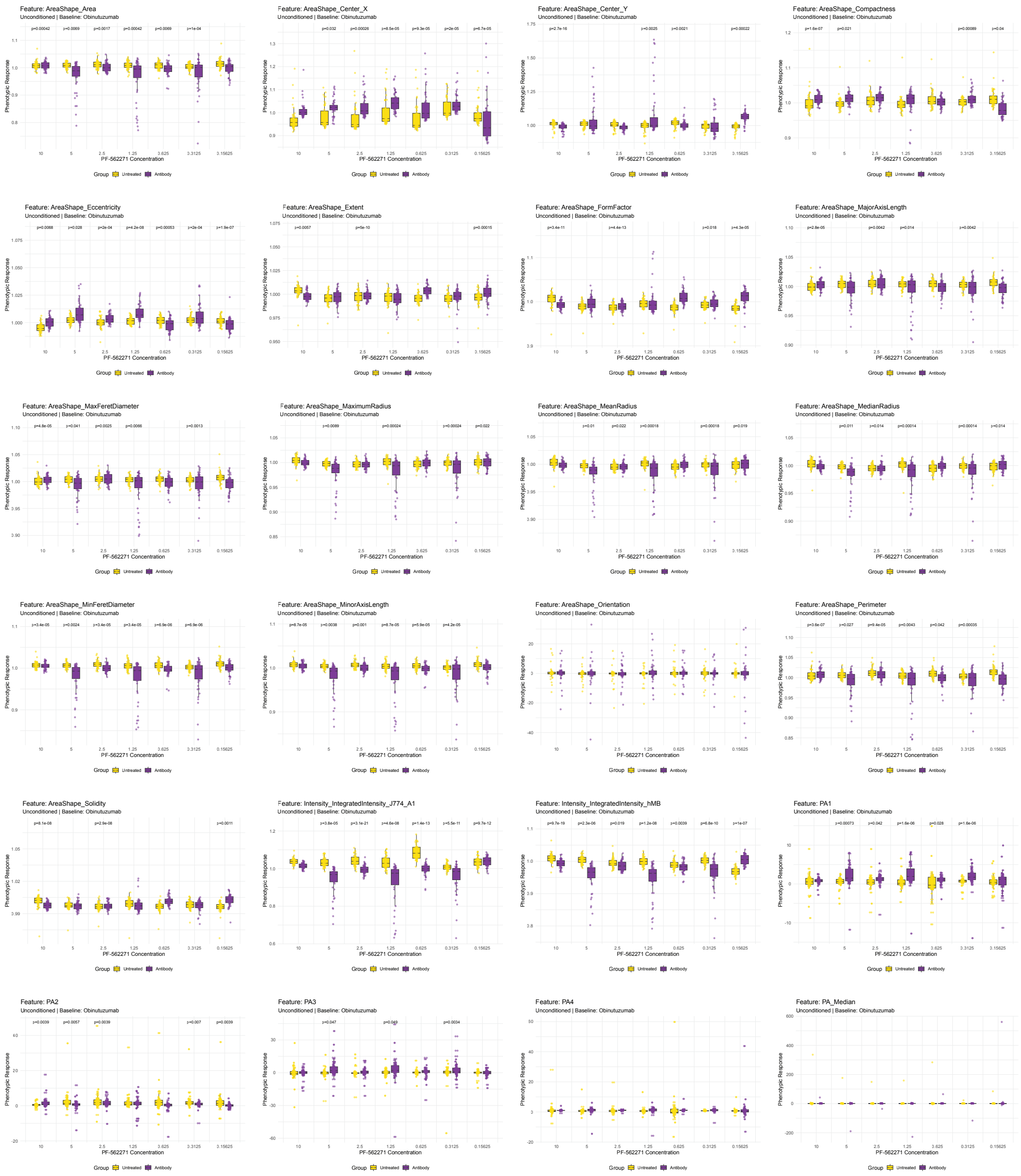

**Supplemental Figure 5: Obinutuzumab timeseries of PF-562271 with per feature phenotypic response calculations with conditioning**

Pooled timeseries per feature Euclidean distance and phenotypic response calculation of all features included in the analysis (n = 24), in the conditioned setting (yellow = PF-562271 monotreatment, purple **Obinutuzumab** + PF-562271 treatment, PF-562271 concentration series [0.15625$\mu$M-10$\mu$M]).

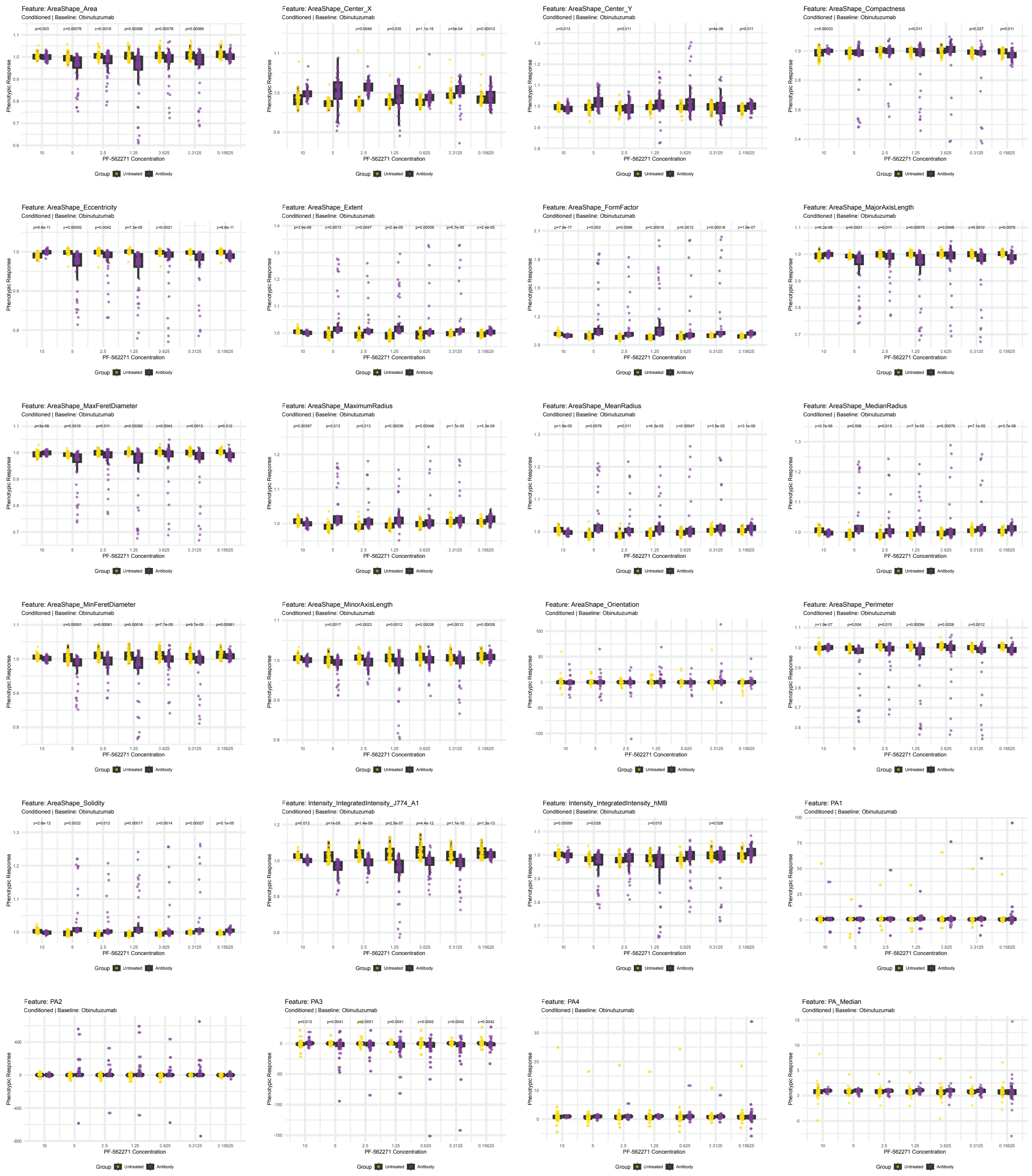

**Supplemental Figure 6: Alemtuzumab timeseries of PF-562271 with per feature phenotypic response calculations without conditioning**

Pooled timeseries per feature Euclidean distance and phenotypic response calculation of all features included in the analysis (n = 24), in the unconditioned setting (yellow = PF-562271 monotreatment, purple Alemtuzumab + PF-562271 treatment, PF-562271 concentration series [0.15625$\mu$M-10$\mu$M]).

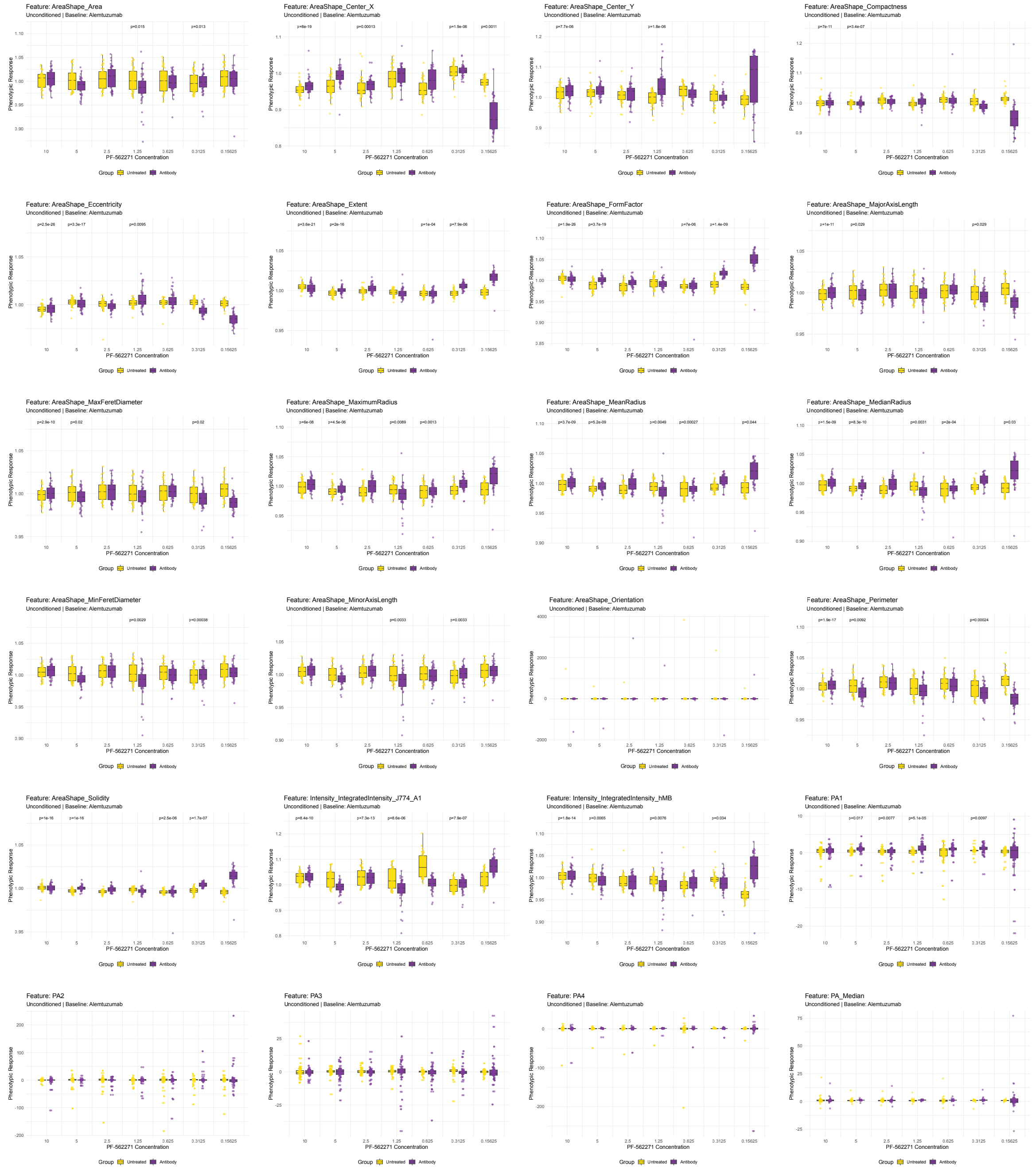

**Supplemental Figure 7: Alemtuzumab timeseries of PF-562271 with per feature phenotypic response calculations with conditioning**

Pooled timeseries per feature Euclidean distance and phenotypic response calculation of all features included in the analysis (n = 24), in the conditioned setting (yellow = PF-562271 monotreatment, purple Alemtuzumab + PF-562271 treatment, PF-562271 concentration series [0.15625$\mu$M-10$\mu$M]).

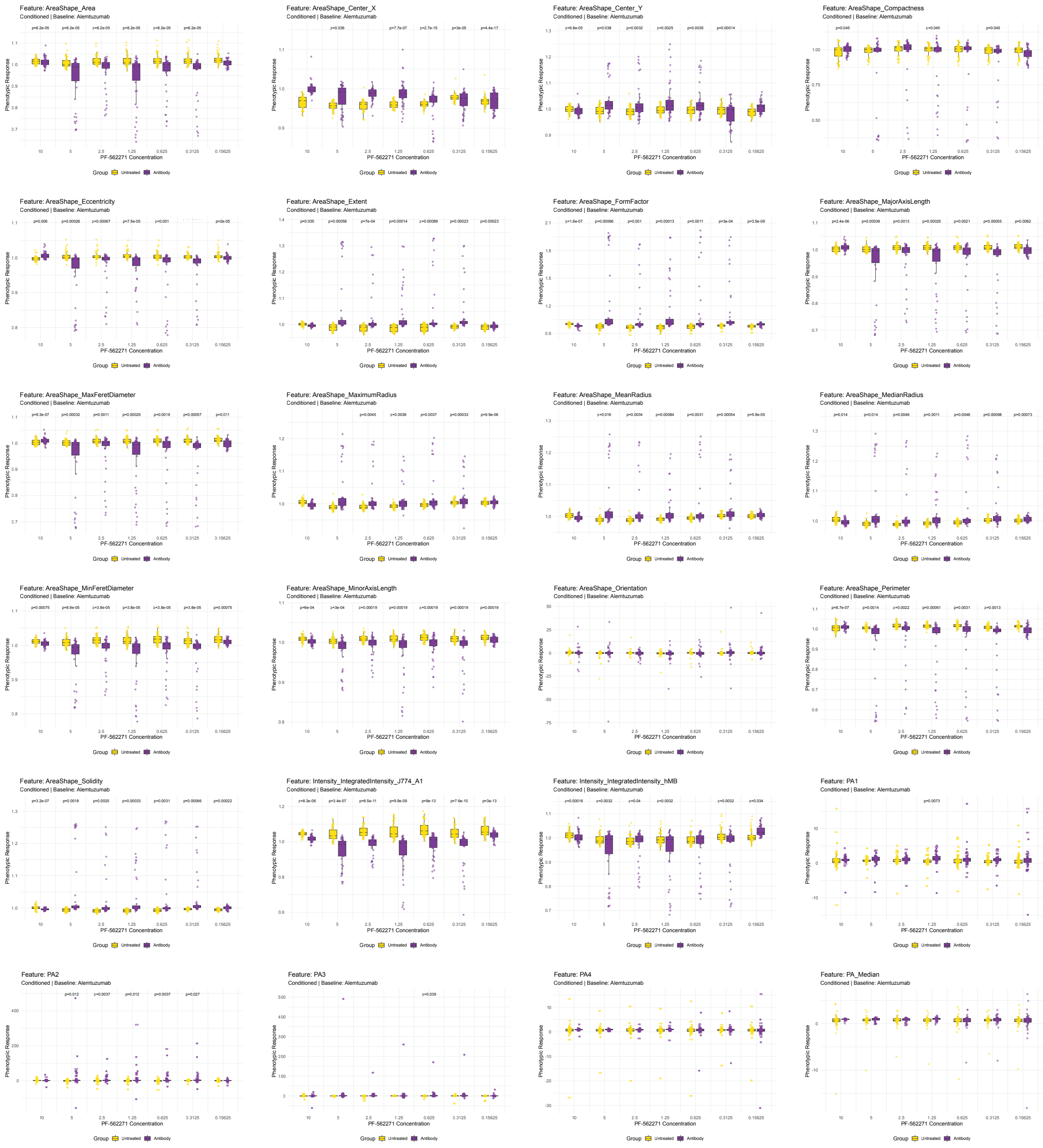

**Supplemental Figure 8: Rituximab limited timeseries of PF-562271 concentration series antibody-dependent cellular phagocytosis by flow cytometry**

Bar graphs showing the ADCP of Rituximab either alone or in combination with PF-562271 concentrations series for either 16 hours (left) or 40 hours (right), and either with (lower) or without conditioning (upper) (grey = Rituximab monotreatment, purple Ritixumab + PF-562271 concentration series [0.15625$\mu$M-10$\mu$M]). Each column represents 1 biological replicate, with row one representing unconditioned 16 hours, row two conditioned 16 hours, row three unconditioned 40 hours, and row four conditioned 40 hours, respectively.

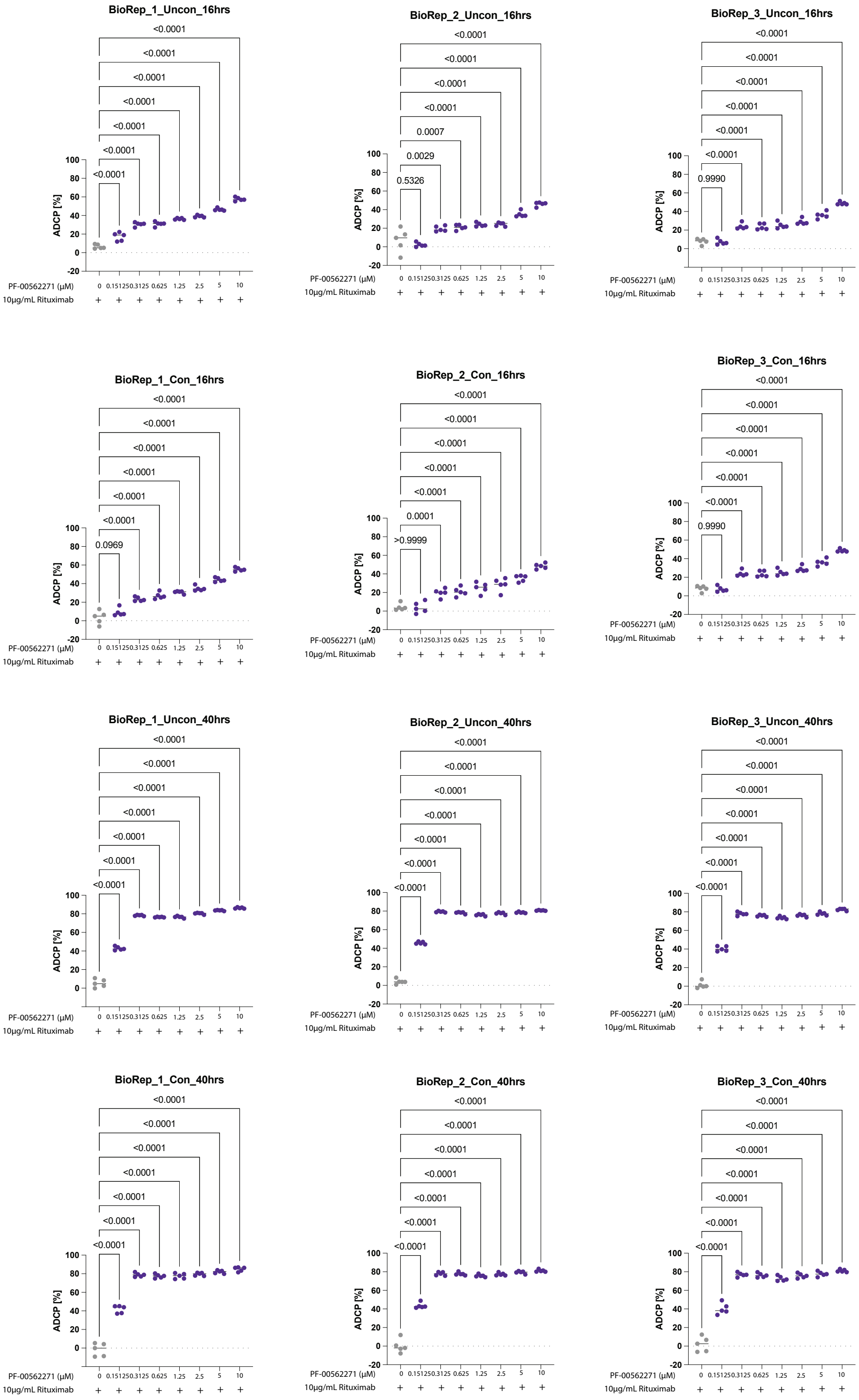

**Supplemental Figure 9: Principal Factor Analysis of CD68+ macrophages under Rituximab and PF-562271treatment *ex vivo***

**A** Elbow plot representing the number of principal factors to include for downstream analysis.
**B** Radar plots showing the variability in area shape features within included principal factors.

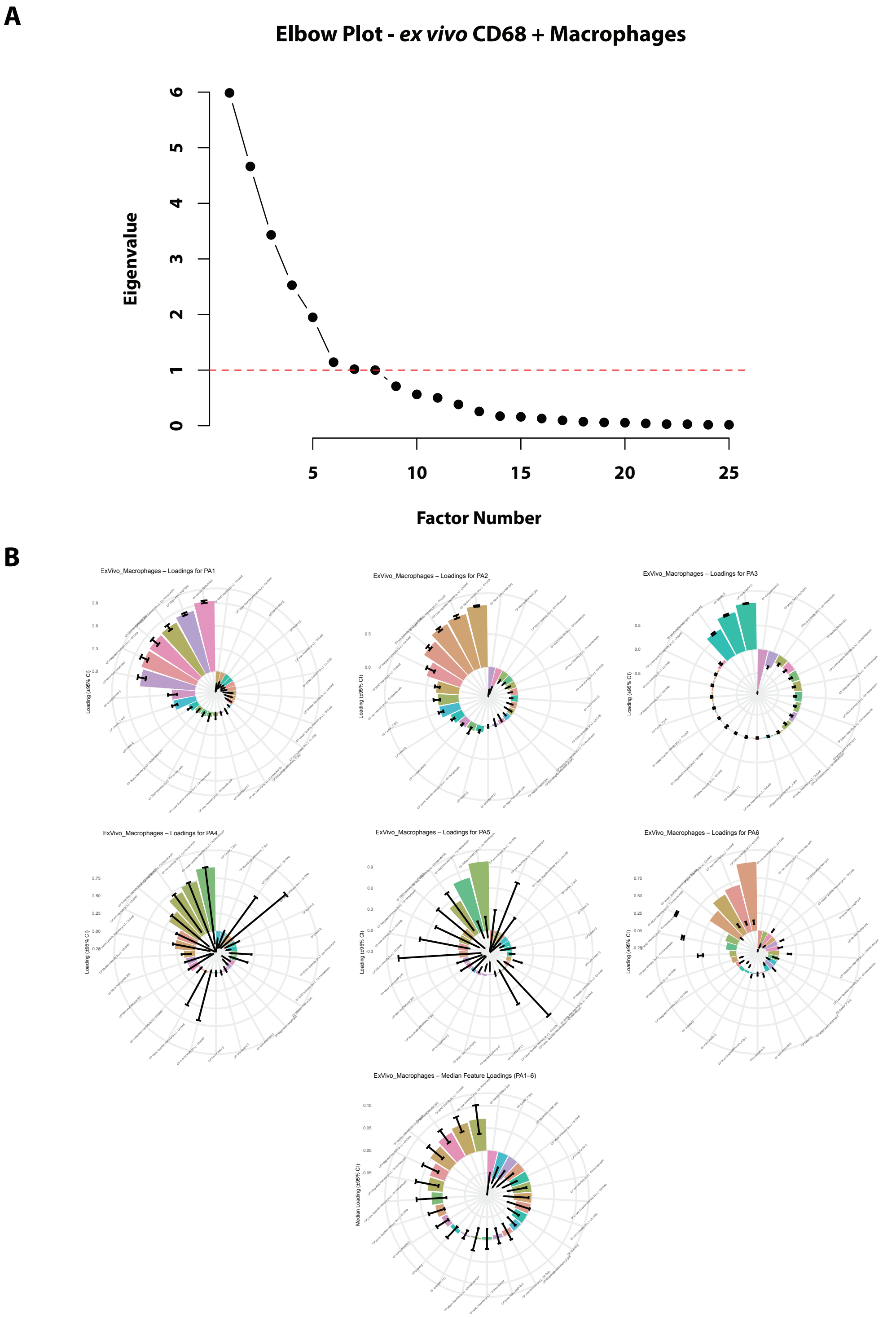

**Supplemental Figure 10: Rituximab and PF-562271 treatment with per feature phenotypic response calculations across all mice**

Pooled mice per feature Euclidean distance and phenotypic response calculation of all features included in the analysis (n = 34), in the conditioned setting (yellow = PF-562271 monotreatment, purple Alemtuzumab + PF-562271 treatment.

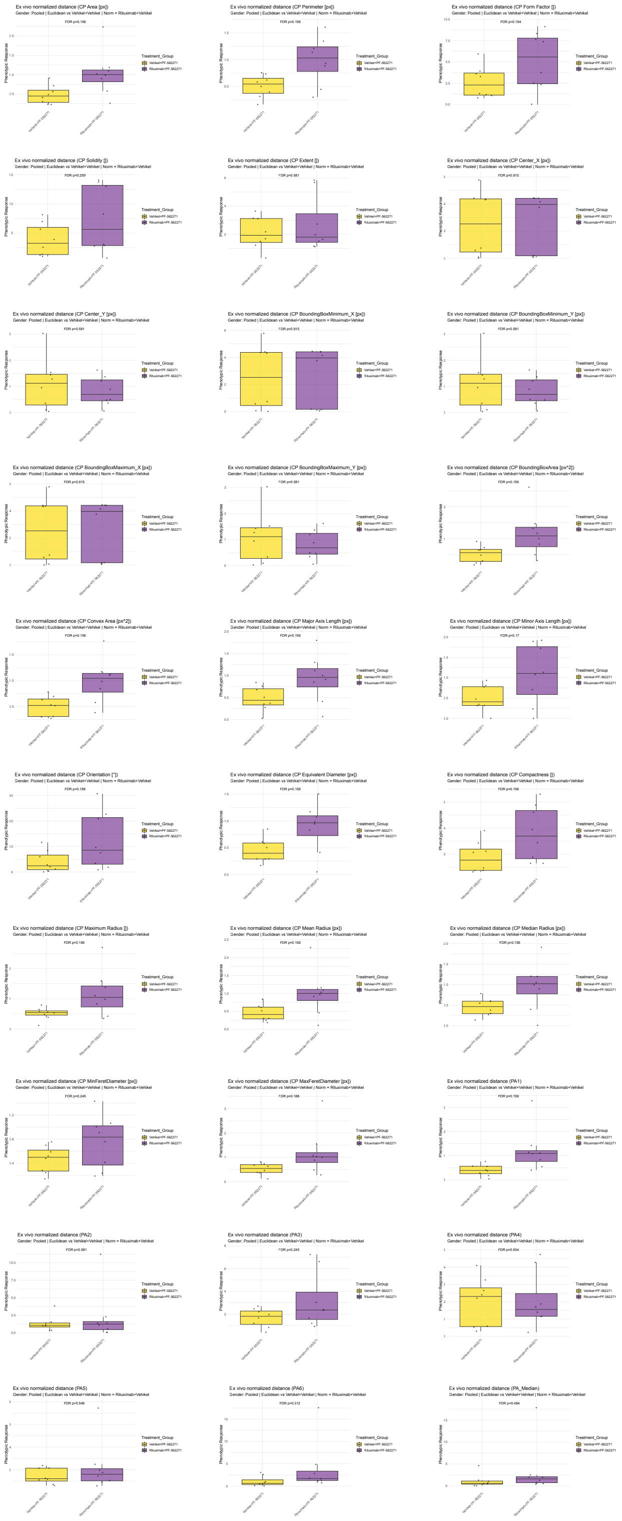

**Supplemental Figure 11: Rituximab and PF-562271 treatment with per feature phenotypic response calculations stratified by male mice**

Male mice per feature Euclidean distance and phenotypic response calculation of all features included in the analysis (n = 20), in the conditioned setting (yellow = PF-562271 monotreatment, purple Alemtuzumab + PF-562271 treatment.

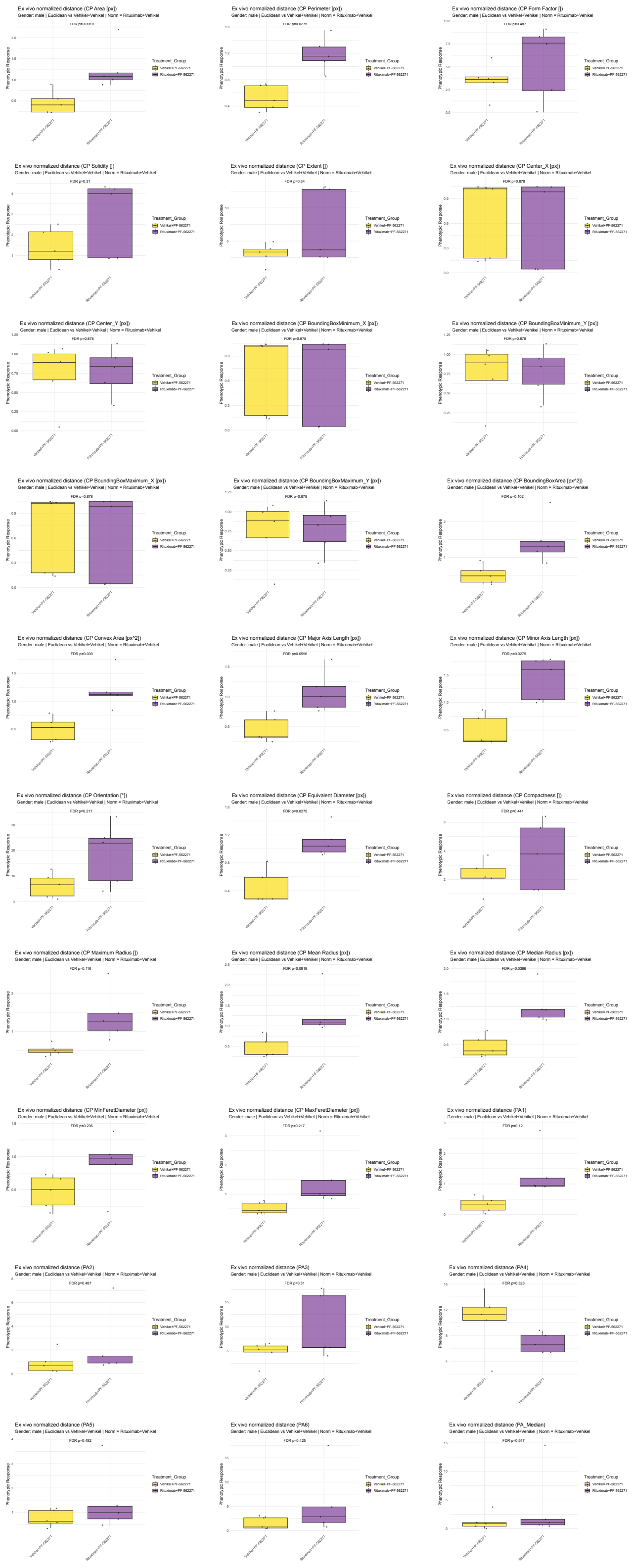

**Supplemental Figure 12: Rituximab and PF-562271 treatment with per feature phenotypic response calculations stratified by female mice**

Female mice per feature Euclidean distance and phenotypic response calculation of all features included in the analysis (n = 14), in the conditioned setting (yellow = PF-562271 monotreatment, purple Alemtuzumab + PF-562271 treatment.

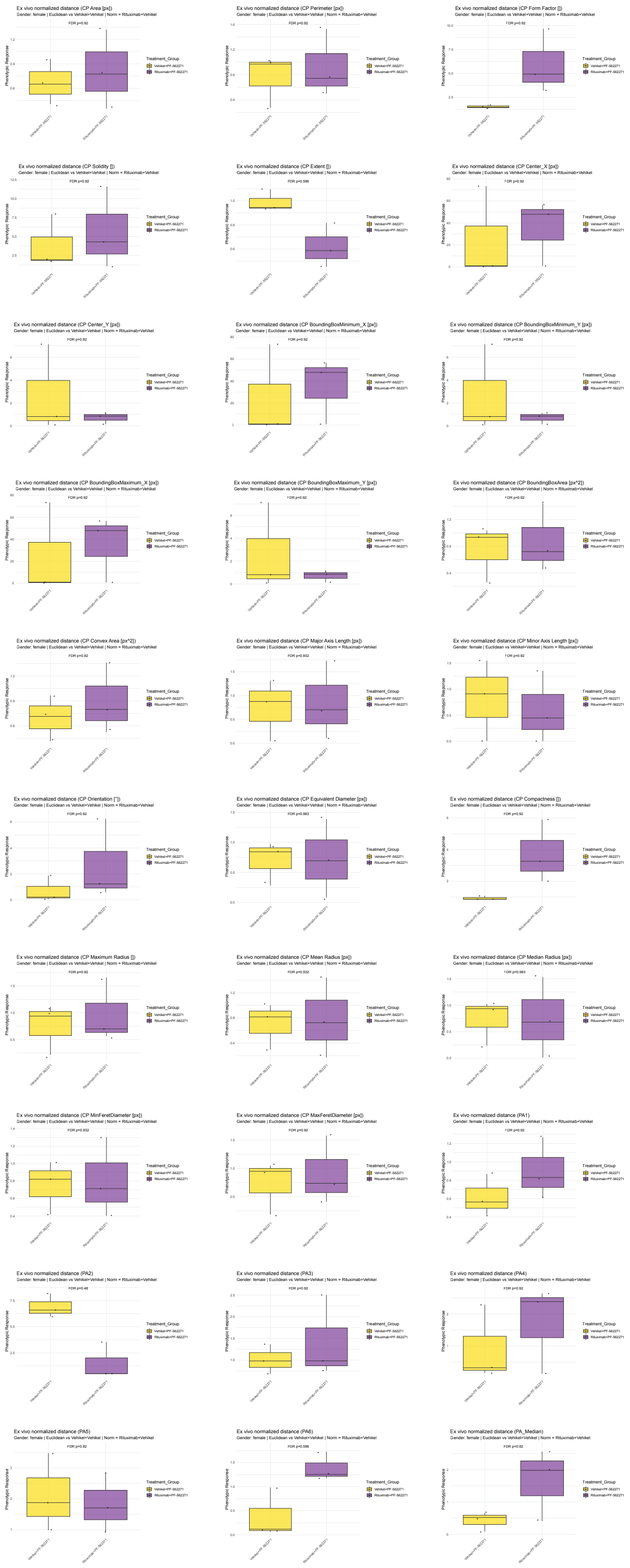

**Supplemental Figure 13: Kaplan Meier survival stratified by gender under Rituximab and PF-562271 treatment *in vivo***

**A** Kaplan Meier post-treatment survival curve of male mice in the cohort (black = vehicle, grey = Rituximab monotherapy, yellow = PF-562271 monotherapy, purple = Rituximab + PF-562271 therapy). **B** Kaplan Meier post-treatment survival curve of male mice in the cohort (black = vehicle, grey = Rituximab monotherapy, yellow = PF-562271 monotherapy, purple = Rituximab + PF-562271 therapy).

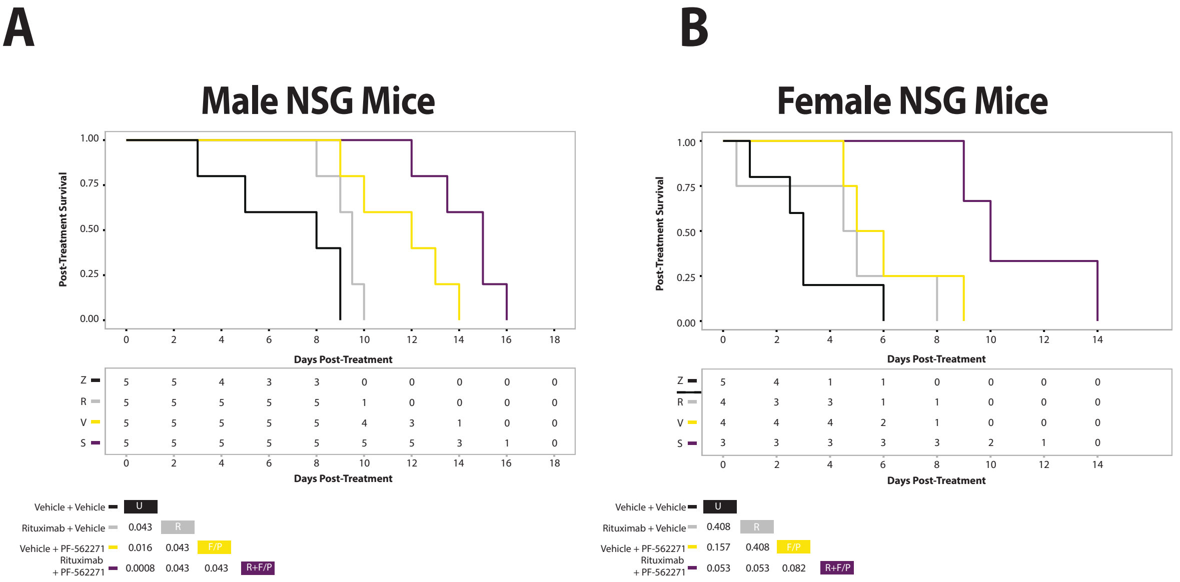
